# Timed chromatin invasion during mitosis governs prototype foamy virus integration site selection and infectivity

**DOI:** 10.1101/2025.01.10.632130

**Authors:** F. Lagadec, P. K. Singh, C. Calmels, D. Lapaillerie, D. Lindemann, V. Parissi, P. Cherepanov, A. N. Engelman, P. Lesbats

## Abstract

Selection of a suitable chromatin environment during retroviral integration is a tightly regulated and multilayered process that involves interplay between viral and host factors. However, whether intrinsic chromatin dynamics during mitosis modulate retroviral genome invasion is currently poorly described. Direct interaction between the spumaretrovirus prototype foamy virus (PFV) Gag protein and cellular chromatin has been described as a major determinant for integration site selection. A previous Gag chromatin-binding site (CBS)–nucleosome co-crystal structure revealed an interaction with the histone H2A-H2B acidic patch via a highly conserved arginine anchor residue. Yet, the molecular mechanisms regulating Gag-chromatin capture during PFV infection remain obscure. Here, we investigated the kinetics of Gag-chromatin interactions during mitosis and proviral integration of PFV-infected synchronized cells. Using Gag CBS variant viruses, we showed that alteration of Gag affinity for nucleosome binding induced untimely chromatin tethering during mitosis, decreased infectivity and redistributed viral integration sites to markers associated with late replication timing of host chromosomes. Mutant Gag proteins were moreover defective in their ability to displace the histone H4 tail from the nucleosome acidic patch of highly condensed mitotic chromatin. These data indicate that the mitotic chromatin landscape during Gag–nucleosome interactions hosts PFV integration site selection determinants and that spumaretroviruses evolved high-affinity chromatin binding to overcome early mitosis chromatin condensation for optimal viral DNA tethering, integration and infection.

## Introduction

Spumaretroviruses, also known as foamy viruses (FVs), are ancient retroviruses marked by a long co-evolutionary relationship with their hosts ^1^. They are prevalent in most non-human primates (NHPs), serving as a reservoir for potential zoonotic transmissions to humans exposed to infected NHPs ^2,3^. FVs establish a persistent, lifelong infection in their human hosts. To date, no reports of severe illness or human-to-human transmission have been documented, indicating effective control of FV replication and transmission in humans^4,5^. The best studied human isolate is called prototype foamy virus (PFV), which constitutes an attractive platform for gene therapy vector development^6^. Additionally, studies of PFV intasomes revealed important structural mechanisms of retroviral integration and inhibition of integrase (IN) activity by strand transfer inhibitors ^7,8^.

Integration of reverse-transcribed viral DNA into the host cell genome is a mandatory step of the retroviral replication cycle^9^. The outcome of the integrated provirus is intricately linked to the local chromatin environment dictating the expression pattern of the viral genome^10,11^. Consequently, retroviruses have evolved genus-specific strategies to navigate within chromosomal features and select suitable chromatin landscapes^12^. Several retroviruses access cellular chromatin during mitosis when the nuclear membrane has dissipated. The impact of mitotic chromosomal condensation dynamics on the process of retroviral integration is however poorly documented. As opposed to the highly pathogenic lentivirus HIV-1, FVs appear to disfavor integration into gene dense regions of the genome but rather target heterochromatin regions such as lamina-associated domains (LADs)^13,14^. The mechanism of integration site selection has been extensively studied for HIV-1^15–18^, uncovering hierarchical interactions between the Gag capsid and IN proteins with cognate cellular factors ^19–24^. In the case of FVs, precise molecular mechanisms responsible for integration site selection remain comparatively obscure and warrant further investigation to inform the applicability of FV-derived gene therapy vectors.

Unlike the case for HIV-1 and other Orthoretroviruses, spumaretroviral Gag is minimally processed by the viral protease, thus sidestepping the production of individualized Gag processing products such as capsid and nucleocapsid^1^. Previously, we demonstrated a direct interaction of PFV Gag with the H2A-H2B acidic patch of host nucleosomes^14^. This interaction critically determined PFV integration targeting, as the R540Q arginine anchor substitution massively redirected integration to centromeric regions of chromosomes. Building from these results, we have further characterized the molecular determinants regulating PFV Gag interaction with chromatin. In addition to the previously-described Arg540 anchor residue, we identified a conserved tyrosine (Y537) that contacts the histone core. Substitution of Gln (Y537Q) altered the ability of PFV Gag to bind nucleosomes *in vitro* and revealed unexpected, untimely chromatin-tethering during mitosis. These phenotypes were moreover associated with a unique pattern of integration site selection, establishing a relationship between mitotic phases, DNA replication-timing and PFV integration site selection.

## Results

### Invariant PFV Gag chromatin binding-site (CBS) residues that interact with human nucleosomes

The crystal structure of PFV Gag CBS (residues 535-550) bound to a nucleosome revealed the molecular determinants for chromatin tethering^14^. The peptide adopts an extended conformation spanning across the histone side of the nucleosome disk (Fig. 1A). One major contact involves PFV Gag residue Arg540 projecting into the H2A-H2B acidic patch cluster formed by H2A Glu61, Asp90 and Glu92 residues (Fig. 1A right panel). This arginine residue is referred to as arginine anchor motif and is a hallmark of many chromatin-binding factors^25^. Amino acid sequence alignment^26^ of CBS regions from several FV species as well as endogenous foamy virus elements revealed a strong evolutionary conservation of this arginine anchor motif (Fig. 1B), suggesting strong selective pressure to maintain Gag chromatin tethering. Further inspection revealed that, in addition to Arg540, the invariant Tyr537 residue interacts with the H2A-H2B acidic patch, with its hydroxyl group making a hydrogen bond with the side chain of H2B residue Gln44. The PFV arginine anchor motif interaction with H2A-H2B acidic patch was shown to be crucial for chromatin tethering and integration site selection^14^. Identification of another conserved acidic patch-interacting residue prompted us to investigate the role of Tyr537 in Gag-chromatin binding and PFV integration.

**Figure 1.**
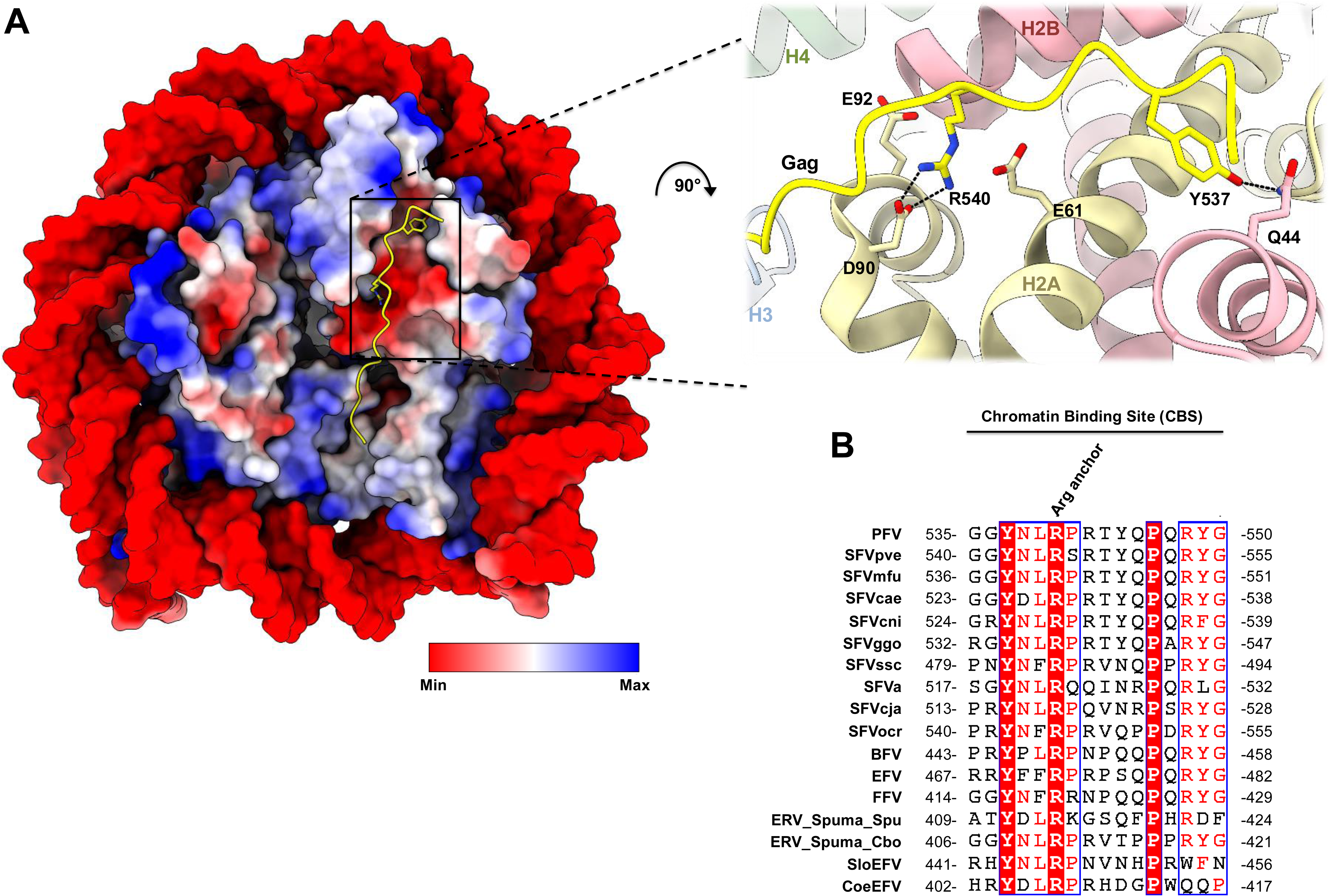
PFV Gag chromatin-binding site (CBS) interaction with a human nucleosome. (A) Overview of the PFV Gag CBS-nucleosome complex structure shown as a surface representation colored as electrostatic potential (left; Protein Database accession code 5MLU). Cartoon representation of the acidic patch engaged by PFV Gag CBS (right). PFV Gag CBS peptide is colored in yellow, histones H2A, H2B, H3, and H4 are shown in pale yellow, red, blue, and green, respectively. (B) Amino acid sequence alignment of Gag CBSs. PFV: prototype foamy virus; SFV: simian foamy virus; pve, *Pan troglodytes verus*; mfu, *Macaca fuscata*; cae, *Chlorocebus aethiops*; cni, *Cercopithecus nictitans*; ggo, *Gorilla gorilla gorilla*; ssc, *Saimiri sciureus*; a, *Ateles* species; cja, *Callithrix jacchus*; ocr, *Otolemur crassicaudatus*; BFV, bovine foamy virus; EFV, equine foamy virus; FFV, feline foamy virus; ERV Spuma Spu, Endogenous retrovirus Spuma *Sphenodon punctatus*; Cbo, *Ciconia boyciana*; SloEFV, Sloth endogenous foamy virus; CoeEFV, Coelacanth endogenous foamy virus. The alignment was performed using ESPript 3

Peptides derived from PFV Gag CBS recapitulated the full-length protein interaction with recombinant nucleosomes^14^. To characterize the role of Tyr537 in nucleosome binding, we performed pull-down experiments using either wild type (WT), Tyr537Gln (Y537Q) or Arg540Gln (R540Q) substitutions in the context of biotinylated CBS peptides. As shown in Fig. 2A, WT CBS peptide efficiently pulled down nucleosomes across tested salt concentrations (75-150 mM NaCl). While the R540Q substitution reduced binding under hypotonic conditions, increasing salt to physiological concentrations (125-150 mM) virtually eliminated binding, which is consistent with the electrostatic nature of the acidic patch interaction (Fig. 1A). The Y537Q substitution also affected nucleosome interaction, though largely in a salt-independent manner. To obtain quantitative data on the interactions between PFV Gag CBS peptides and nucleosomes, we performed bio-layer interferometry (BLI). Streptavidin biosensors were loaded with either WT, R540Q or Y537Q biotinylated CBS peptides and probed for interaction with various concentrations of recombinant nucleosome core particles (NCPs) at physiological salt concentration. Binding of nucleosomes to Gag CBS peptides induced a wavelength shift that was directly linked to the number of molecules bound to the surface of the biosensor. As expected, based on the results of the pull-down experiments, the strongest BLI interaction was observed with WT CBS peptides (Fig. 2B). For each nucleosome concentration, Y537Q and R540Q CBS peptides showed ∼5 fold and almost complete reduction of the interaction with nucleosomes, respectively. These data show that while PFV Gag arginine anchor motif at position 540 is a major determinant for NPC binding, the invariant Tyr537 residue supports and enhances the Gag-nucleosome interaction.

**Figure 2.**
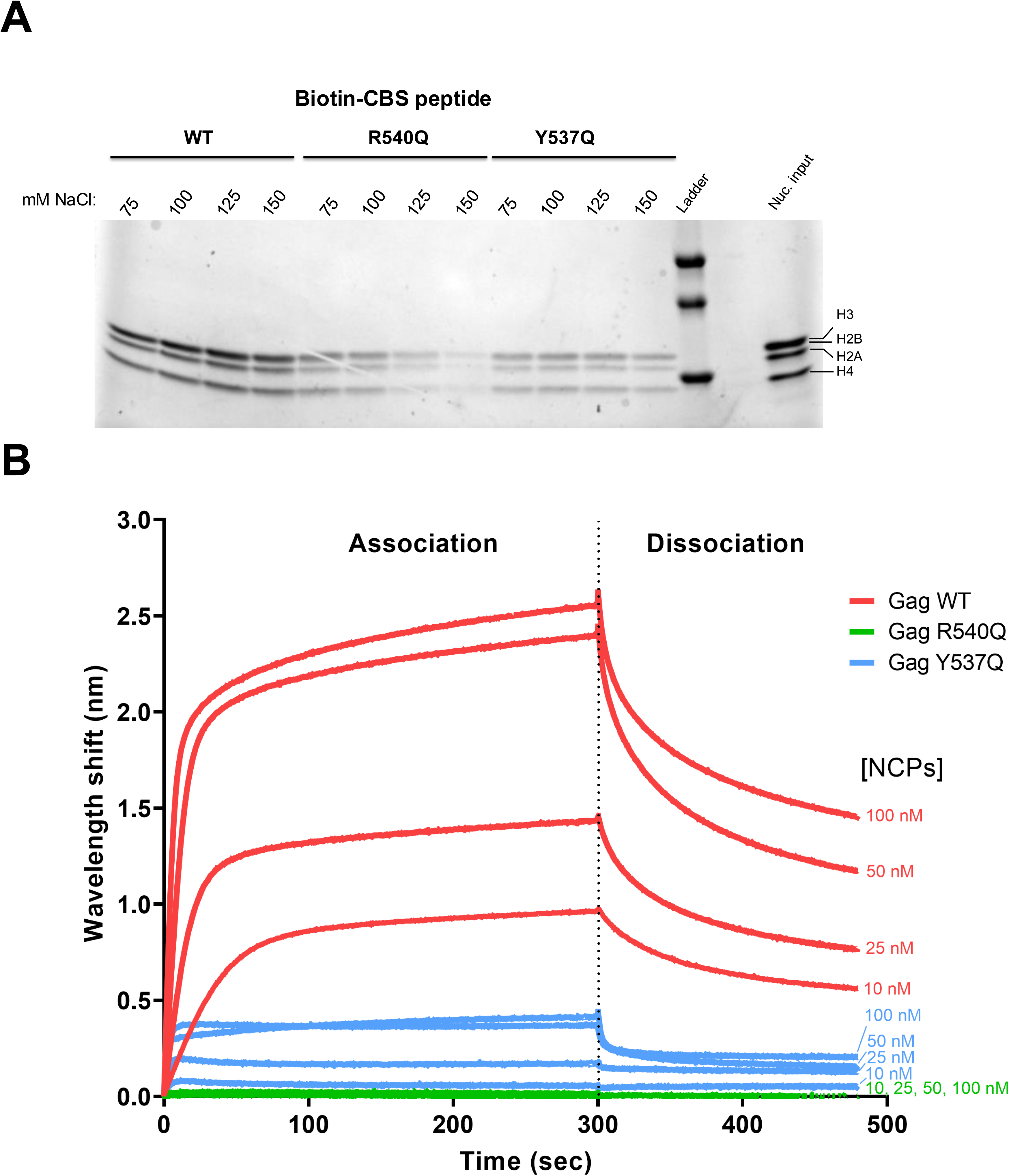
PFV Gag residue Y537 contributes significantly to the interaction with nucleosomes. (A) Streptavidin pull-down of recombinant nucleosomes with biotinylated Gag CBS peptides in the presence of 75-150 mM NaCl. (B) Bio-Layer interferometry (BLI) sensorgram of immobilized Gag CBS peptides and different concentrations of recombinant NCP. The binding intensity (nm) is normalized with a buffer condition without NCP. Results are representative of 3 independent experiments.

### Invariant CBS residues Tyr537 and Arg540 influence the timing of chromatin capture during mitosis

Mitosis is required during foamy virus infection to gain access to host chromatin^27^. Upon cell division, nuclear envelope breakdown exposes chromatin to incoming FV particles for subsequent integration of the viral genome. To evaluate the precise role of conserved CBS residues in chromatin invasion during the course of mitosis, we infected G2/M-synchronized HT1080 cells with single-round PFV particles encoding Gag WT, R540Q or Y537Q. Cells were arrested at the G2/M boundary using cyclin-dependent kinase 1 (CDK1) inhibitor RO-3306^28^. Removal of the molecule by washing allowed the arrested cells to progress into mitosis in a synchronous fashion (Fig. S1 and S2). To synchronize PFV infection, virus particles were added to G2/M-arrested cells 1 h prior to RO-3306 removal, which induced Gag accumulation at the microtubule organizing complex (MTOC)^29^ (Fig. 3). Infected cells were fixed at different time points post G2/M block release, representing different mitotic phases, and stained by immunofluorescence using anti-Gag and anti-Lamin A/C antibodies. Lamin A/C staining was used as a marker for nuclear envelope integrity and chromatin accessibility. The percentage of Gag chromatin occupancy was quantified at each mitotic phase (Fig. 3 and Fig. S3). Staining of incoming WT Gag particles showed that following centrosome duplication in prophase, chromatin tethering was concomitant with nuclear envelope breakdown at late prophase/early pro-metaphase (Fig. 3, top panels). The viral proteins decorated cellular chromosomes during further mitotic steps and the nuclei of newly divided interphasic cells. Consistent with previous reports, although PFV Gag R540Q particles accumulated at the MTOC, they were effectively excluded from chromatin at all stages of mitosis and interphasic cells^14,30,31^ (Fig. 3, middle panels). PFV Gag Y537Q particles adopted a unique phenotype. Indeed, from the beginning of mitosis and until metaphase, Gag Y537Q phenocopied Gag R540Q behavior, with MTOC retention and exclusion from chromatin. However, Y537Q particles shifted from this chromatin-excluded state to a chromatin-bound state during late mitosis, at telophase (Fig. 3, bottom panels). This unexpected result revealed that despite chromatin juxtaposition after nuclear envelope breakdown, an additional layer of regulation prevented Gag Y537Q–chromatin interactions until the late stages of mitosis. These data revealed that in addition to altering nucleosome binding, substitutions of conserved CBS residues can induce a delay in chromatin capture during mitosis.

**Figure 3.**
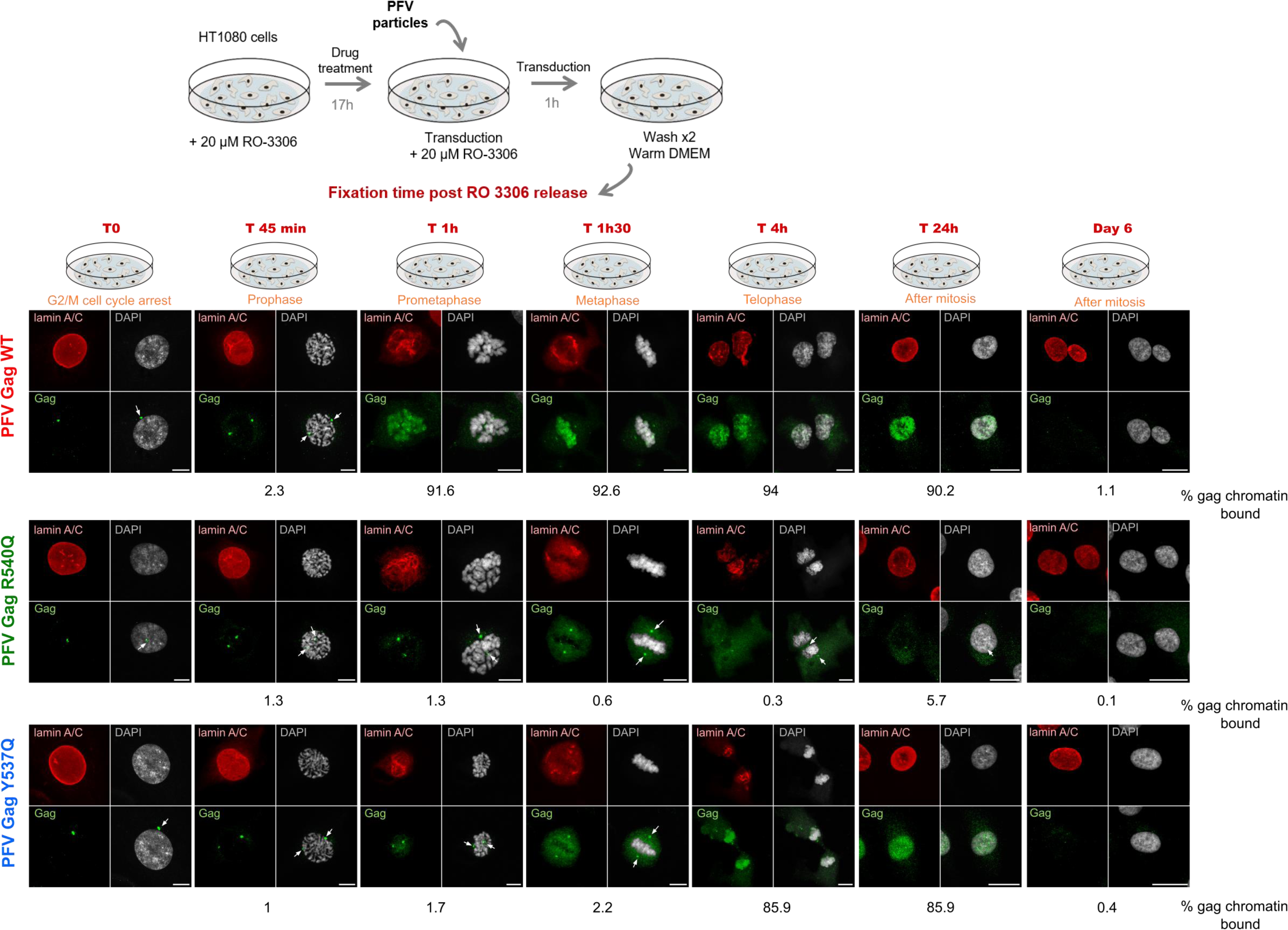
Conserved Gag CBS residues are essential for timely mitotic chromatin capture. Gag localization during PFV infection. G2/M-phase synchronized HT1080 cells were transduced with PFV particles encoding WT, R540Q or Y537Q Gag and fixed at different time points post drug release, corresponding to different mitotic phases indicated in orange. Gag proteins were detected using polyclonal anti-PFV Gag antiserum (green); the nuclear envelope was stained with anti-lamin A/C antibodies (red) and cellular DNA with DAPI (grey). White arrows show MTOC accumulation of PFV Gag. Scale bars: 20 µm. Percentages of chromatin-bound Gag are indicated for each mitotic phase. Results are representative of those observed across at least 5 independent experiments.

### Untimely capture of mitotic chromatin influences PFV fitness

We next investigated the effect of invariant CBS residue substitutions on viral infectivity. WT and CBS mutant PFV particles were purified by ultracentrifugation through 20% sucrose cushions and the quantity of particles used for each transduction was normalized by immunoblotting and real-time PCR for relative viral genome copy number determination (Fig. S4A). For an infection-defective control, we used IN active site mutant virus IN-NQ. HT1080 cells were transduced with equal amounts of viral particles and cultured for six days before being subjected to flow cytometry to quantify the percentage of GFP-positive cells. As expected, IN-NQ virus was unable to establish a productive infection (Fig. 4A). Consistent with previous results, R540Q led to a ∼40% decrease of GFP positive cells (Fig. 4A)^14,31^. Gag Y537Q mutant virus showed a more moderate, but still significant, ∼25% decrease of infectivity.

**Figure 4.**
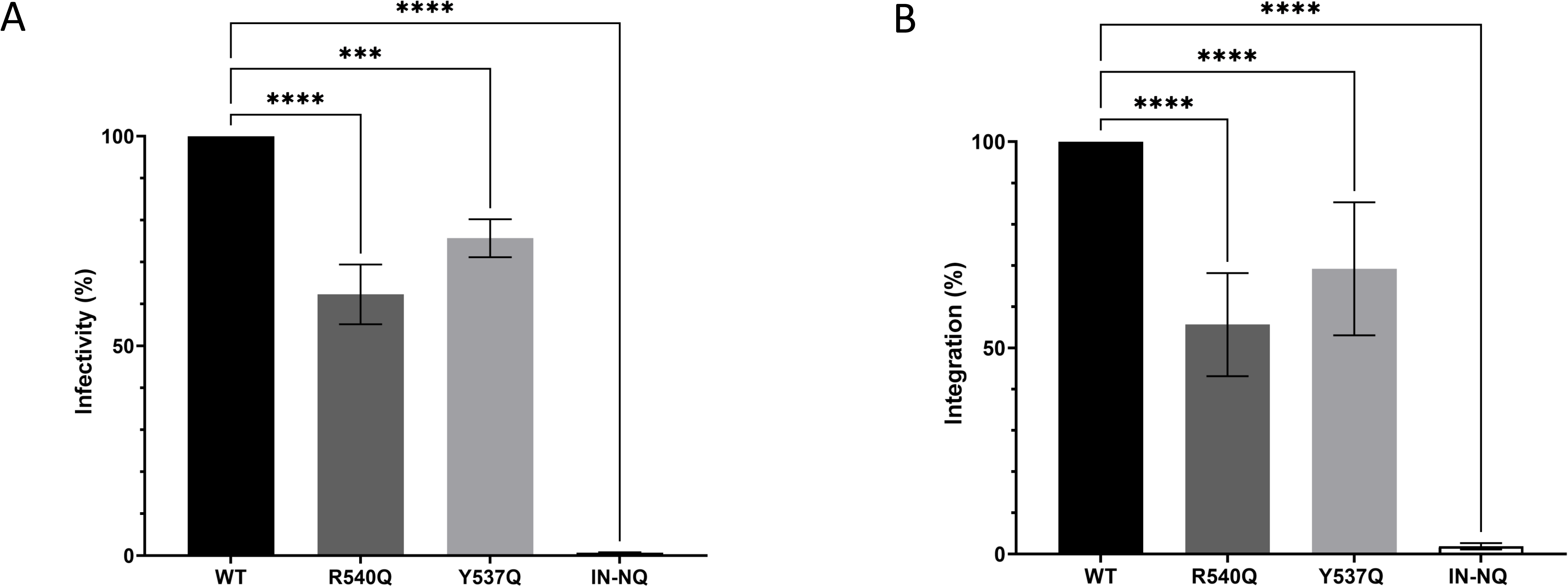
Conserved PFV Gag CBS residues are required for optimal infectivity. (A) Six days post-infection, GFP positive cells were counted by flow cytometry as relative measures of infectivity. Error bars are SDs determined from at least three independent infections; the WT values in each experiment were set to 100%. (B) Quantitative PCR of integrated vDNA, six days post HT1080 infection, with PFV vector particles carrying WT, R540Q or Y537Q Gag with WT IN or WT Gag containing virus with catalytically inert D185N/ E221Q IN (IN-NQ). Results are expressed as percentage relative to the WT condition, which was set to 100%. Statistical analysis were performed using the ordinary one way ANOVA, withTukey’s multiple comparisons tests *** p < 0.0005; **** p < 0.0001).

As particles containing Gag Y537Q showed delayed chromatin tethering compared to WT (around 3 h, Fig. 3 and Fig. S3), we next measured whether this effected the relative timing of integration. To this end, we synchronously infected HT1080 cells with WT, R540Q or Y537Q particles and blocked the integration step at different times post-infection using the IN inhibitor dolutegravir (DTG). DTG was maintained at 2 µM in the cell growth medium until flow cytometry analysis 6 days post-infection. If the delayed chromatin tethering observed with CBS mutants delayed integration, the kinetics of infection should be differentially sensitive to DTG treatment. As seen in Fig. S4B, the integration kinetics of all three viruses were highly similar. This suggests that the untimely chromatin tethering observed with PFV Gag CBS substitution mutants did not affect the timing of integration.

Next, we performed real-time PCR using primers specific to the PFV long terminal repeat (LTR) region to measure integrated proviral content 6 days post-infection. As observed in Fig. 4B, integration was barely detectable for IN-NQ control virus, and infections performed with PFV Gag R540Q and Y537Q mutant particles showed ∼40% and ∼35% decreases from the WT, respectively. These data reveal an important role for early chromatin tethering by PFV Gag during mitosis to ensure optimal genome capture for subsequent integration.

### Y537Q redirects integration to late-replication markers of host chromatin

Our previous results showed that the R540Q substitution led to massive redistribution of integration events into centromeric regions of the genome^14^. We next investigated the effect of the Y537Q variant on integration site selection, comparing the results to previously determined WT and R540Q sites^14^ as well as a reference set of *in silico-*selected random integration control (RIC) sites (Table S1). Sites of Y537Q integration revealed a unique pattern of proviral distribution (Fig. 5 and Table S2). Although maintaining a preference for centromeric regions versus the WT, Y537Q targeted centromeres less robustly than did R540Q (P=0.035; Table S2). By contrast, Y537Q showed the greatest preference for Giemsa-positive cytobands (gp100) and lamina-associated domains (LADs), and was least biased for elements associated with active chromatin including gene-dense regions, transcriptional activity and speckle-associated domains (SPADs). While WT PFV integration was enriched near transcriptional start sites (TSSs) and CpG islands, these features were avoided by the Y537Q and R540Q viruses.

**Figure 5.**
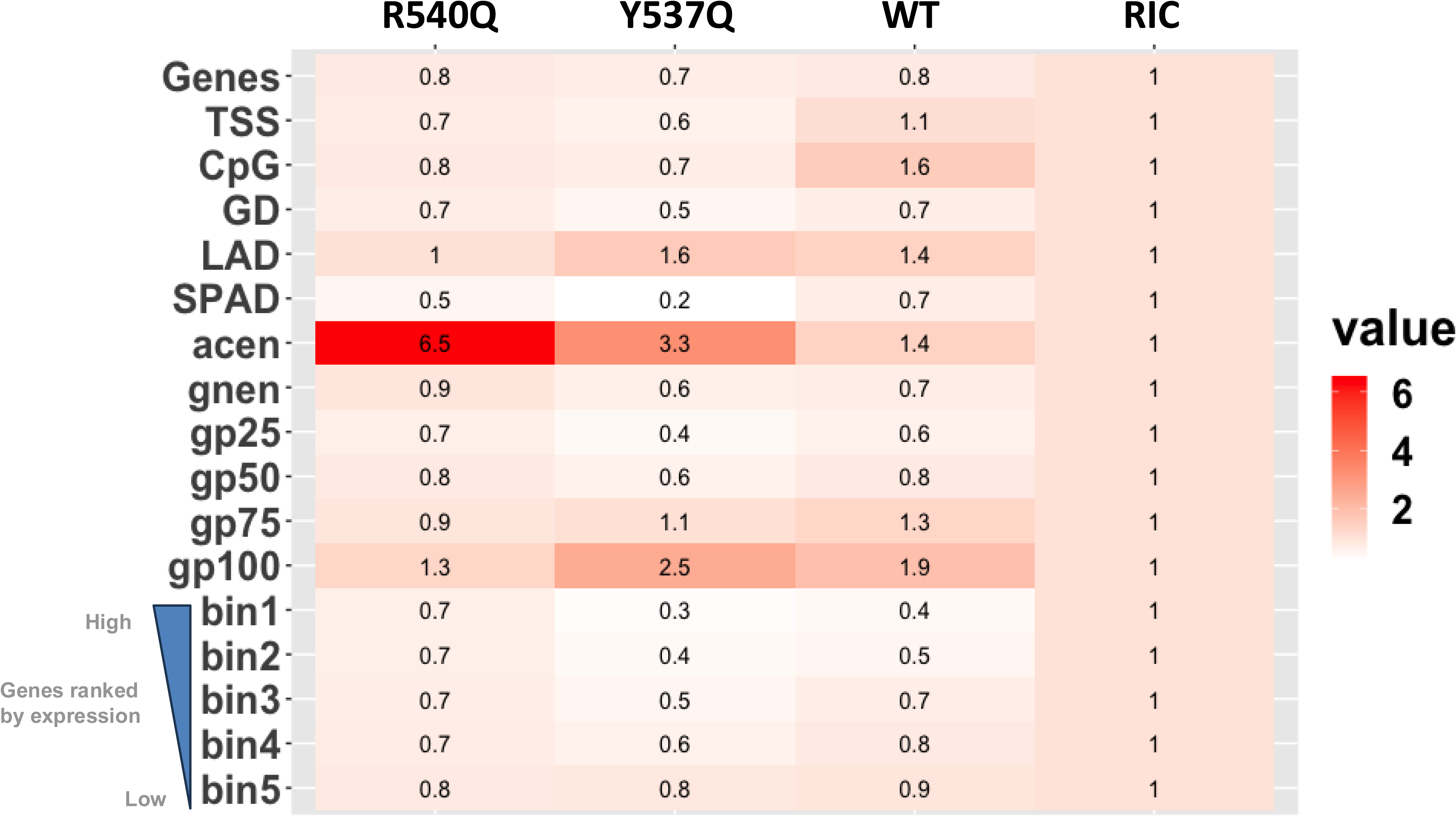
Integration site distributions of WT and Gag CBS mutant viruses. Integration frequencies normalized to in silico-calculated random integration controls (RICs) are shown as a heatmap. Values > 1 (red color) indicate enrichment of PFV sites compared to random, whereas values < 1 (white color) represent features avoided by PFV for integration. Genes were divided into five groups based on expression, with bin1 being top-expressed genes. Human cytobands specific to genome build hg38 are shown as acen, gnen, gp25, gp50, gp75 and gp100. TSS, GD, LAD, and SPAD represent transcription start site, gene density (+/- 0.5 Mb), lamina-associated domain, and speckle-associated domain..

Compared to the R540Q mutant, Y537Q integration disfavored gene-dense regions of chromatin (P=0.009; Table S2). To assess the granularity of this difference, we plotted the percentage of integration sites per Mb on each autosomal chromosome, which revealed a significant preference for Y537Q to integrate into a subset of human chromosomes (Fig. 6A). Chromosomes 21, 4, 18, 13 and 6 were among the most frequent targets of Y537Q integration (Table S3 and S4). Conversely, chromosomes 22, 19, 17, 15, 16, and 20 were highly avoided for integration. These chromosome clusters differ with respect to replication timing, as chromosomes 22, 20, 19, 17, 16 and 15 are the earliest replicating chromosomes while chromosomes 21, 18, 13 and 4 are comparatively late replicating^32^. To further probe this connection, we stratified chromosomes into three groups based on replication timing, and plotted percent integration per Mb (Fig. 6B). WT and the Gag CBS mutants generally preferred late-replicating as compared to early-replicating chromosomes for integration. However, compared to R540Q, Y537Q targeted early replicating chromosomes significantly less (P<10^-10^) and late replicating chromosomes significantly more (P<10^-10^) (Fig. 6B and Table S5). Despite the difference in integration for various genomic features such as centromeres (Fig. 5), R540Q and WT PFV, moreover, similarly targeted early and late replicating chromosomes for integration (P=0.6 and 0.9 for early and late replicating chromosomes, respectively). To independently validate this observation, we correlated chromosomal distributions of replication-initiation determinant protein (RepID) occupancy^33^ with PFV integration site distributions. As seen in Fig. 6C, chromosomes 22, 19, 17 and 20 were highly enriched for RepID whereas chromosomes 21, 13, 4 and 18 were comparatively RepID-poor. Consistently, we observed a significant inverse correlation between RepID enriched-chromosomes and Y537Q integration sites (R^2^=0.66; R^2^ for WT and R540Q=0.28 and 0.12, respectively) (Fig. 6D, Table S6). The seven chromosomes least enriched for RepID were most frequently targeted by Y537Q, whereas RepID-enriched chromosomes were less targeted by Y537Q compared to WT and R540Q, confirming the integration bias of Y537Q for late replicating chromosomes (Fig. 6C and D).

**Figure 6.**
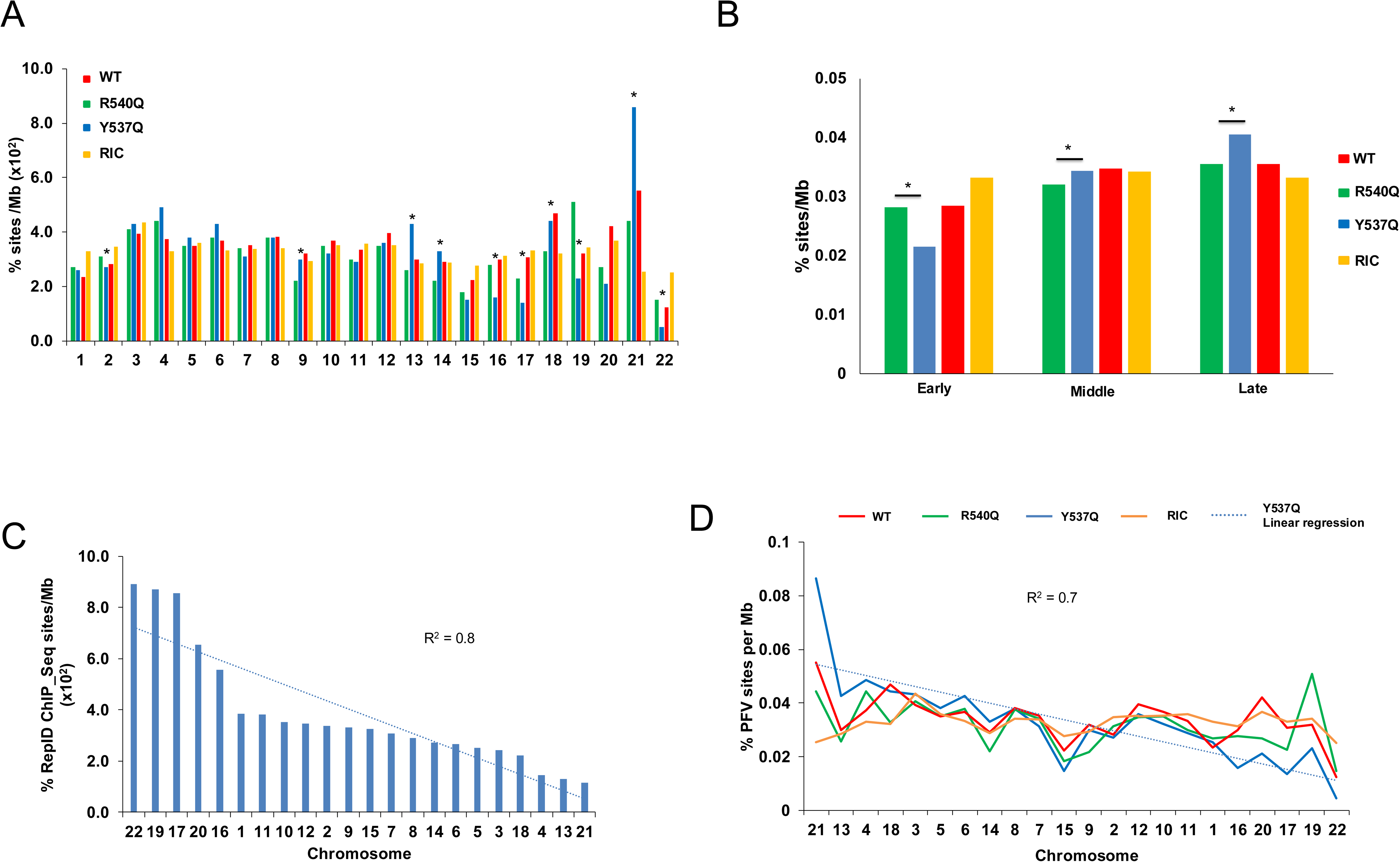
Chromosomal distributions of PFV proviruses. (A) PFV integration sites % per Mb (Y-axis) of indicated human chromosomes (X-axis) are shown for R540Q and Y537Q viruses along with WT PFV and random integration control (RIC). Significant differences in chromosomal targeting between R540Q and Y537Q is shown (Fisher’s exact test; p < 0.05). (B) PFV integration % per Mb for three chromosomal groups based on replication timing: early, middle, and late replicating (X-axis). PFV integration per Mb for each group is shown (* p < 0.05; Fisher’s exact test). (C) Chromosomal distribution of replication-initiation determinant protein (RepID) ChIP-Seq sites per Mb. (D) Correlation between chromosomal distribution of RepID binding sites per Mb and % PFV integration sites per Mb.

We next correlated RepID occupancy across human genes, which revealed that genes devoid of RepID sites accounted for the overall similar preference for Y537Q and R540Q to target genes for integration (Fig. 5) (P=0.08) (Fig. S5C and Table S7). In contrast, RepID-associated genes were significantly less targeted by Y537Q for integration (P<10^-10^) (Fig. S5B and Table S7). Compared to the mutants, WT PFV differentially targeted RepID-associated genes, yet targeted genes devoid of RepID sites similarly as the CBS Gag mutants for integration (Fig. S5B-C). Consistent with the reduced preference for Y537Q to target early-replicating chromosomes for integration, Y537Q integrated significantly less frequently into ±2.5 kb regions of RepID sites than did WT and R540Q (P<10^-10^) (Fig. S5A and Table S7).

We next sought to independently verify the unique pattern of Y537Q integration targeting. The human genome can be represented as 10 Spatial Position Inference of the Nuclear genome (SPIN) states^34^, which are radially distributed from the nuclear center (Speckles SPIN state) to the periphery (Lamina state). Correlating WT, R540Q, and Y537Q PFV integration sites with SPIN states revealed that all three viruses disfavored integration within the seven innermost states (Speckles, Act1, Act2, Act3, Repr1, Repr2, and Lm1), while the three outermost SPIN states (Lm2, Lamina_Like, and Lamina) were significantly more targeted than random (Fig. S6A and Table S8). Notably, Y537Q integration positively correlated with SPIN states from the center to the periphery (R^2^=0.6) while WT (R^2^=0.3) and R540Q (R^2^=0.2) exhibited integration patterns more similar to the RIC (R^2^=0.3) (Fig. S6). These findings indicate that the Y537Q PFV mutant shifts integration from central gene-dense genomic regions to more peripheral regions of the genome. Since LADs, which represent late-replicating genomic regions, are predominantly in the Lamina state, these findings support the results presented in Fig. 6.

### Differential gene targeting by PFV Gag CBS mutant viruses

Because Y537Q Gag binding to mitotic chromatin was significantly delayed compared to the WT (Fig. 3 and Fig. S3), we next investigated the connection between mitotic chromatin dynamics and PFV integration. To examine chromatin state transitions, we analyzed published data on mitotic-specific transcription^35^, which reported gene expression levels at different time points (t=0, 40, 80, 105, 165, and 300 min) after the release of nocodazole-induced mitotic arrest. We compared the expression of genes at t=105 min to their expression at three earlier time points, t=0, 40, and 80 min. Genes were identified as mitotic-specific at t=0 (early), t=40 (middle), and t=80 (late mitosis) if their expression at these time points was at least 1.5-fold higher than at t=105 min. These mitotic-specific genes were then ranked by transcriptional activity and then halved based on expression level. Using these six gene sets, we assessed their enrichment for early-, middle-, or late-replicating chromosomes (referred to as early-, middle-, and late-replicating genes) in top50% vs. bottom50% genes at each time point. This analysis revealed that well-expressed genes at t=0 and t=40 min were significantly enriched for early replication (P<0.00001; Fisher’s exact test) (Table S9). However, by t=80 min (late mitosis), this enrichment was lost (P=0.08). Additionally, we observed that the bottom half of expressed genes were enriched for late replication at t=0, t=40 and t=80 min (P= <0.01; Fisher’s exact test). These findings indicate that early-replicating genes are preferentially expressed at t=0 and t=40 min, while late-replicating genes are underrepresented in top50% genes. As mitosis progresses, these enrichment patterns are lost, with early- and late-replicating genes becoming more randomly distributed (or distributed as per their expected genome frequencies) between highly and less-expressed gene groups.

Our results demonstrated that the Y537Q mutant targets early-replicating chromosomes significantly less than the WT and R540Q mutant, while preferentially targeting late-replicating chromosomes (Fig. 6B). Moreover, Y537Q and R540Q mutants differentially targeted RepID-associated genes (early-replicating) versus non-associated genes (late-replicating) (Fig. S5B and S5C). Given the distinct targeting patterns of these mutants with respect to replication timing, we next investigated their preferences for transcriptional activity. Because the upper 50% of expressed genes at t=0 and t=40 min were predominantly early-replicating (Table S9), we hypothesized that these genes would be differentially targeted by Y537Q versus R540Q for integration, while the lower half of expressed genes would be similarly targeted. As expected, compared to R540Q, Y537Q disfavored integration into the top 50% of expressed-genes (Fig. 7A-C; Table S10). Reciprocally, both mutants similarly targeted comparatively poorly expressed genes across time points (Fig. 7D-F).

**Figure 7.**
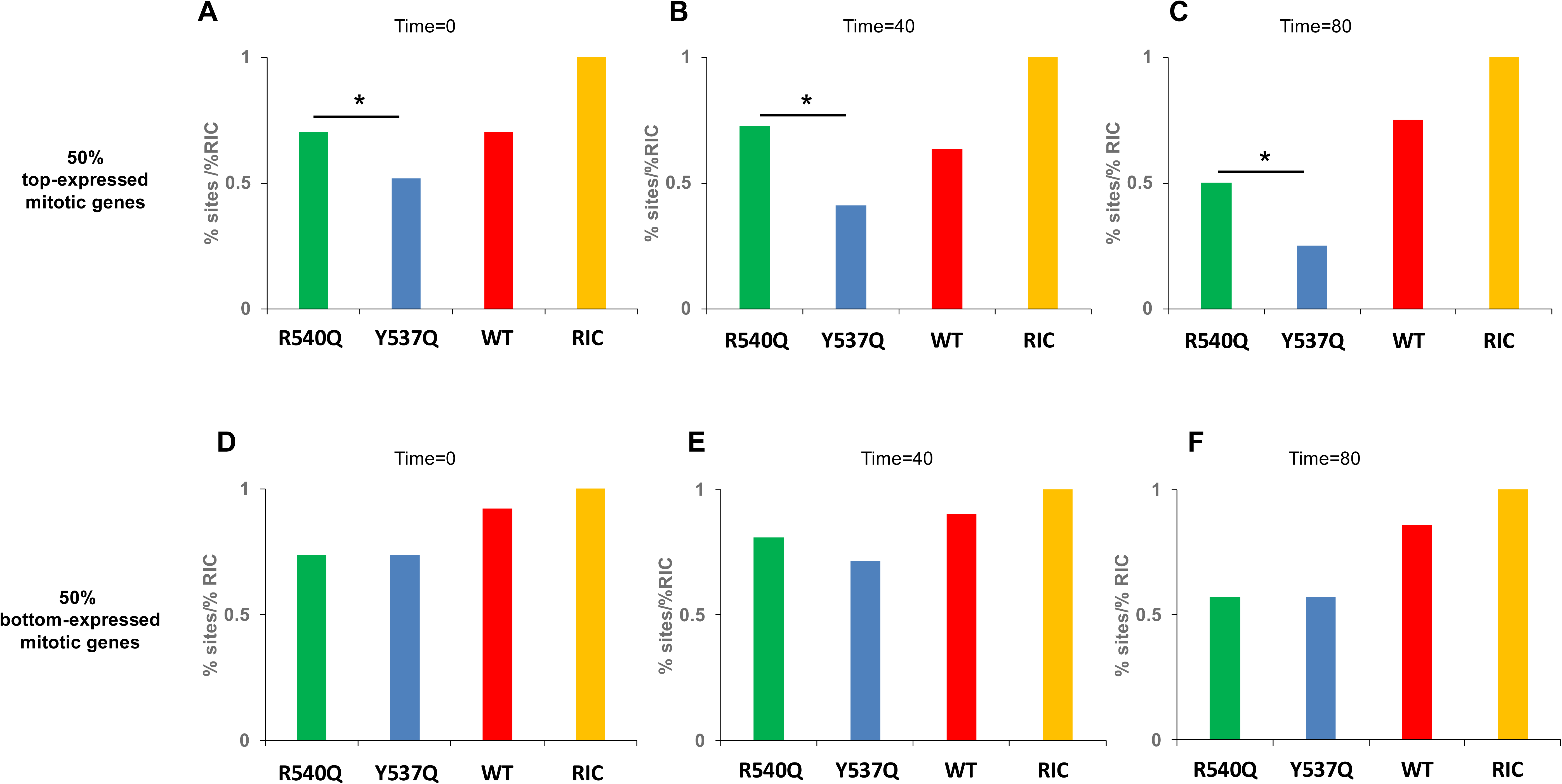
PFV integration sites within mitotic expressed genes. Mitotic genes were classified at 0 (A and D), 40 (B and E), and 80 (C and F) min if the expression of the gene was ≥1.5 times its expression at 105 min from release of mitotic arrest. PFV genic sites (%) are shown for 50% top-expressed (A, B and C) and 50% bottom-expressed mitotic genes (D, E and F). *, significant difference in targeting between R540Q and Y537Q (Fisher’s exact test; p < 0.05).

### WT PFV Gag but not CBS mutant Y537Q displaces the H4 tail from the acidic patch

The phenotype observed for Y537Q chromatin engagement during mitosis (Fig. 3 bottom panel) prompted us to investigate the mechanisms underlying its delayed access. During mitosis, chromatin condensation increases until the end of metaphase^36^. In late anaphase and telophase, mitotic chromosomes decondense to re-establish interphase chromatin^36^. The H2A-H2B acidic patch Gag docking station is an important determinant for chromatin compaction by hosting the neighboring nucleosome amino-terminal H4 tail^37–40^. In such condensed early mitotic chromatin, nucleosome acidic patches are preferentially occupied by H4 tails. To explain the Gag Y537Q phenotype, we hypothesized that depending on H2A-H2B acidic patch accessibility during the course of mitosis, chromatin segues from a condensed refractory state to a more open, binding-permissive state. In such a scenario, WT PFV Gag has the ability to compete for the H4 tail–acidic patch interaction, explaining its early chromatin binding mode during virus infection, while Gag Y537Q requires more opened chromatin with comparatively accessible acidic patches. To test our hypothesis, we performed BLI competition experiments using immobilized PFV Gag CBS peptides challenged with NCP saturated with H4 tails to mimic occupied acidic patches from condensed early mitotic chromatin. We first monitored binding of WT H4 tail (H4 WT) versus nucleosome-interaction deficient mutant (H4 mut) peptides to purified NCPs. WT H4 tails, but not the mutated versions, efficiently interacted with recombinant NCPs (Fig. S7). Next, we probed PFV Gag peptide capacity to compete with the H4 tail for H2A-H2B acidic patch binding. As seen in Fig. 8A, WT PFV Gag CBS interaction kinetics with NCPs under saturating concentration of WT H4 tails, but not the H4 mut tails, revealed a modified, slower association curve. Under these competition conditions, the interaction reached the same level as without H4 tails, and were consistent with displacement events in favor of the PFV Gag CBS. The same experiment was performed with immobilized PFV Gag R540Q (Fig. 8B), which recapitulated the results observed in Fig. 2B, namely no interaction with the NCP. With CBS peptide Y537Q, the results in Fig. 8C showed that PFV Gag Y537Q was unable to effectively compete for H2A-H2B acidic patch binding with nucleosome saturated by H4 WT or H4 mut (as seen in Fig. S7, H4 mut peptide retains some residual binding that, in excess, is sufficient to prevent Gag Y537Q binding). These BLI experiments revealed that WT PFV Gag CBS peptides retain chromatin binding capacity under conditions of excess H2A-H2B acidic patch interacting competitor (H4 tail). Conversely, binding capacity of PFV Gag Y537Q CBS peptides for the nucleosomes was drastically reduced by the presence of the H4 tail. These data are compatible with the hypothesis that H4-occupied acidic patches in condensed mitotic chromatin restrict Gag Y537Q binding.

**Figure 8.**
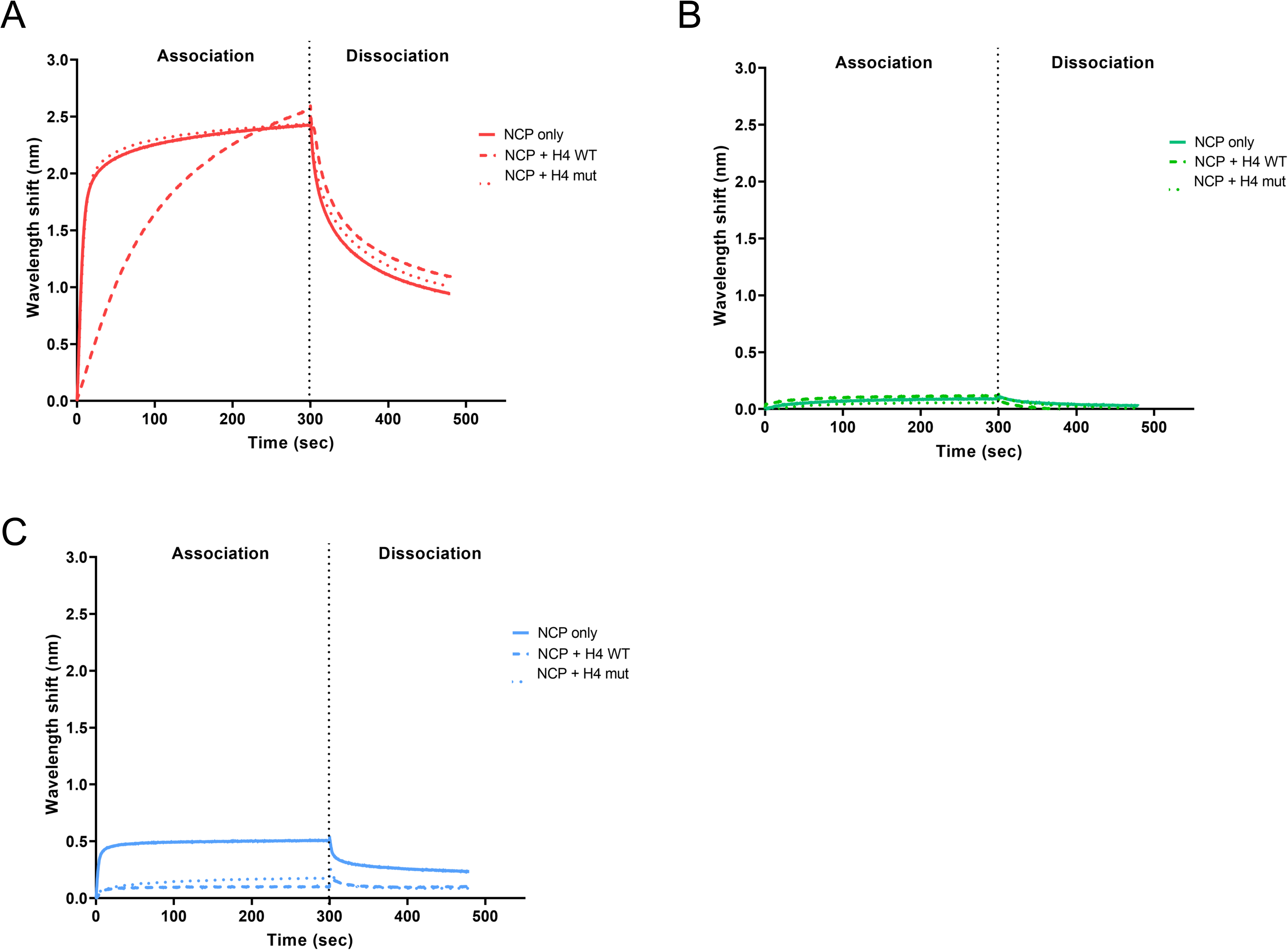
Interaction of the H4 tail to the nucleosome acidic patch prevents Gag Y537Q binding. Bio-Layer interferometry (BLI) sensorgram of free nucleosomes (NCP, plain line), saturated with WT H4 tail (residues 2-24) (dashed line) or H4 mut (triple alanine substitution) (dotted line) binding to immobilized (A) WT Gag (red), (B) R540Q Gag (green) and (C) Y537Q Gag (blue) peptides. Binding intensity (nm) was normalized to conditions without NCP. The mean of 2 independent experiments is plotted.

## Discussion

Here we show that the PFV Gag CBS harbors invariant residues crucial for H2A-H2B acidic patch engagement and integration site selection. In addition to the well described arginine anchor motif R540, we identified a highly conserved Y537 residue supporting the interaction with nucleosomes. Although not essential for binding with histones, substituting this tyrosine for glutamine decreased NCP capture ∼5-fold. These results are consistent with previous observations^41,42^ and highlight the crucial role of CBS Gag residues in interacting with the nucleosome acidic patch. Nucleosome binding is a prerequisite for many cellular factors to fulfill chromatin transactions. A recent comprehensive study showed that more than 50% of nucleosome interactions are mediated by the acidic patch^43^. The plethora of nucleosome complex structures solved by cryo-EM provided an extensive source of molecular information governing acidic patch engagement. They revealed a strong conservation of the arginine anchor motif but also illuminate the variety of secondary contacts within the acidic patch^25,44^. As seen with Gag Y537Q, such secondary contacts may tune the affinity requirements for chromatin interactions.

In contrast to lentiviruses, spumaviruses cannot infect resting cells and require nuclear envelope breakdown during mitosis to access chromatin^27^. We tracked PFV Gag proteins after infecting cell-cycle synchronized cells with viral particles harboring CBS amino acid substitutions R540Q or Y537Q. WT particles showed practically immediate chromatin tethering upon nuclear envelope breakdown. Altering nucleosome binding with the Y537Q substitution unexpectedly delayed chromosome capture until late mitosis (telophase). We provided biochemical evidence that Gag Y537Q’s incapacity to bind early mitotic chromosomes is likely due to its inability to displace the H4 tail from condensed chromatin acidic patches. Indeed, several reports showed a major role for the H4 tail – acidic patch interaction in high order chromatin structure^45,46^. Additionally, disruption of this interaction, such as in the case of pioneer transcription factors, leads to a destabilization of chromatin packing, facilitating its opening for subsequent DNA transactions^47^. It will be of interest to determine whether PFV Gag possesses the capacity to alter local chromatin structure by evicting the H4 tail from the acidic patch and the potential role on viral and cellular functions^44^.

Accessing and persisting in the nuclear compartment is a challenge for many viruses. The nuclear envelope represents an impassable barrier for several viruses that rely on mitosis to access chromatin^48^. Here we confirmed the important role of PFV Gag tethering to chromatin. Indeed, affecting Gag’s capacity to engage chromatin provoked a modest but significant decrease of viral infectivity, with a decrease of integrated viral DNA. In addition, infections with Gag CBS variants induced a redistribution of integration sites along the host genome with a marked preference of the Gag Y573Q virus to target late-replicating chromosomes (Fig. 6B). Our finding that Y537Q fails to bind chromatin in early mitosis (Fig. 3 and Fig. S3) is consistent with the integration site analysis (Fig. 7), which showed a significant depletion of Y537Q integration into highly expressed genes that are enriched for early-replicating chromosomes. As mitosis progresses, the proportion of mitotic-specific genes from early-, middle-, and late-replicating chromosomes becomes random, and the enrichment of early-replicating genes diminishes (Table S9). This transition suggests an increased accessibility of late-replicating chromosomes to Y537Q Gag binding and subsequent integration, explaining why the Y537Q mutant preferentially targets late-replicating regions of chromatin in late mitosis for integration.

Integration site selection is a common feature of retroviruses although each genus evolved specific mechanisms to select their insertion loci. However, we note that the Y357Q phenotype (Fig. S6A) resembles the behavior of the CPSF6 binding-deficient HIV-1 N74D capsid mutant virus, which shifts HIV-1 integration sites from gene-dense, centrally located SPAD/Speckle-enriched chromosomes to gene-poor, LAD/Lamina-enriched chromosomes^17,18,24^. We furthermore note that a recent study has described SPADs as early-replicating chromosomal regions^49^. Because previous research indicated that speckle depletion did not affect HIV-1 integration into SPADs^50^, it seems that replication timing, in addition to spatial organization of chromosomes, may play a role in lentiviral DNA integration targeting. We plan in the future to further explore the relationship between DNA replication timing and Orthoretroviral DNA integration.

We previously described the PFV Gag CBS as a pan-nucleosome binder important for maintaining integration site distribution^14^. Herein we revealed that the observed integration site selection phenotype of CBS variants is the consequence of delayed chromatin capture and the spatial chromosomal re-organization along the course of mitosis that, depending on the mitotic stage, exposes different chromosomes and genes to PFV integration complexes. In light with the correlation between tethering timing during mitosis and integration selectivity, it is tempting to speculate a potential opportunistic mechanism of spumaretroviral integration site selection; Gag-containing pre-integration complexes will interact with the first suitable and available nucleosomes to dictate eventual integration location. To best succeed, foamy viruses evolved a strong chromatin binding capacity in order to capture highly condensed chromatin early in mitosis. This ensures optimal chromatin retention, integration and viral infectivity. Harnessing this intrinsic, unique chromatin binding capacity of spumaretroviral Gag might constitute a promising starting platform for the development of targeting-tunable FV-based vectors in gene therapy.

## Material and Methods

### Nucleosome core particle (NCP)

Recombinant NCPs (mono-nucleosomes) were assembled as previously described^13^. Briefly, individual human histones H2A, H2B, H3 and H4 were purchased (The Histone Source; Colorado State University) and octamers were assembled by dialysis. The Widom-601 DNA^51^ was generated by restriction enzyme digestion of a concatenated construct and purified by anion exchange chromatography onto a POROS-HQ column. NCPs were assembled with a ratio 1.1 : 1 (1.1 octamer : 1 DNA) by successive steps of salt dialysis against TEN buffer (2 M-0 M NaCl, 10 mM Tris pH 7.5, 1 mM EDTA, 1 mM DTT). NCPs were kept at -80°C in 0 mM NaCl TEN buffer.

### Pull down assay

20 µL of streptavidin beads (Dynabeads™ MyOne™ Streptavidin T1; invitrogen; Thermo Fisher Scientific) were used per condition. StreptaBeads were washed twice with pull down buffer (20 mM Tris HCl pH 7.5, 0.1 % NP 40, 10 % Glycerol, 75 to 150 mM NaCl, 0.5 mM DTT and 1 mM PMSF) containing 75 mM of NaCl and 100 µg of BSA, followed by one wash in the same conditions, without BSA. 1.5 µg of Gag CBS biotinylated peptide and 3 µg of recombinant NCPs were added per condition, in a final volume of 800 µL of pull down buffer with corresponding NaCl concentrations. The reaction mix was incubated 2 h on a rotation wheel, at 4°C. Beads were washed three times with the corresponding NaCl concentration buffer, and bound proteins were eluted by the addition of 10 µL of SDS Sample Buffer 2.5X. Samples were separated by SDS-PAGE, and analyzed after colloidal Coomassie blue staining.

### BLI analysis

Quantitative interaction experiments were performed by Bio-Layer Interferometry (BLI), using a BLItz instrument (Octet ®N1; Sartorius). Two different runs were performed: the first one, into water, to coat biosensors with biotinylated peptides and the second, into Blitz buffer (1X Phosphate Buffer Saline (PBS) pH 7.4, 0.02% (v/v) Tween-20, 0.1% (w/v) bovine serum albumin (BSA)), to fix NCPs on immobilized peptides. All steps were performed under 2200 rpm shaking speed. Streptavidin biosensors (Octet®SA Biosensors; Reference 18-0009; Sartorius) were first pre-wet for at least 10 min with water. The baseline, with only water, was measured for 60 sec. 10 µM of Gag CBS biotinylated peptides (or H4 biotinylated peptides), diluted in water, were loaded onto the biosensors, for 180 s, to reach a binding of ∼1.5 nm, followed by a dissociation step of 180 s. Following the first run, coated biosensors were equilibrated for at least 10 min in Blitz buffer. The second run started with 60 s of baseline, followed by the association step, where biosensors were dipped into serial dilutions of NCPs (10, 25, 50 or 100 nM), for 300 s. Finally, the dissociation step was performed in Blitz buffer, for 300 s. For competition experiments, the second run started with 60 s of baseline in Blitz buffer, followed by 300 s of association step. During this step, solutions of 100 nM of NCPs were pre-incubated with or without 100 µM of H4 peptide (WT or mutant, non-biotinylated), in Blitz buffer, for at least 30 min at room temperature. Finally, the dissociation step was performed in Blitz buffer, for 180 s. Non-specific binding controls were performed, such as binding of 100 nM of NCPs on biosensors without biotinylated peptides loaded, as well as binding of Blitz buffer on loaded biosensors. For each time point measured, data of this second control (binding of Blitz buffer on biosensors loaded with biotinylated Gag WT peptide) were subtracted to every condition and were plotted on curves, using Prism 9 software.

### Cell culture

Human fibrosarcoma cell line HT1080 (ATCC CCL-121) and proteoglycan-deficient packaging cell line 293T-25A^52^ were maintained in Dulbecco’s modified Eagle Medium (DMEM; Fisher), supplemented with 50 µg/mL of gentamicin (Fisher Scientific) and 10% fetal calf serum (FCS) (Eurobio Scientific).

### Viral vectors productions

PFV particles were produced using a four-component system, based on a protocol described in ^52^. Plasmids pcoPG4 (PFV Gag), pcoPP (PFV Pol), pcoSE (SFVmac/SFVmcy Env) and puc2MD9 were co-transfected using a ratio 4:2:1:28. Briefly, 293T-25A cells were transfected using calcium phosphate, in DMEM without FCS, for 5 h at 37°C. After 5 h, transfection medium was replaced by fresh DMEM containing 10% FCS, for 48 h at 37°C. Supernatants were harvested, filtered through a 0.45 µM filter, either kept at -80 °C or processed for viral particles purification. Viral particles were pelleted by ultra-centrifugation of supernatants, 2 h at 25 000 rpm, at 4°C, on a 20% sucrose cushion, in a SW-32Ti rotor. PFV particles were re-suspended in 1X PBS supplemented with 5% DMSO, and kept at -80°C.

### Viral transductions

To analyze kinetics of PFV integration, 5.10^4^ HT1080 cells were platted in each well of a 24 well plate. To remove residual contaminant plasmids, purified viral particles were treated with 1 µL of Denarase, 25 kU (20804-25k, c-LEcta), in a final volume of 200 µL of PBS 1X, for 1 h 30 min at 37°C. Synchronization of transductions was performed using spinoculation, as previously described^53^. Briefly, viral particles were added to cold DMEM, and 500 µL of viral solution were added to each well. The plate was kept 10 min at 4°C and centrifuged 30 min at 1200 g, at 10°C. Infection medium was then replaced with pre-warmed DMEM. This time point was considered as the starting point (T0) of the transduction. For integration kinetic experiments, 2 µM DTG (GSK1349572, Selleckchem) (or DMSO as a control) were added at different time points post-transduction. Transduced cells were maintained for 6 days post-transduction, with fresh medium containing 2 µM DTG replaced every two-days. 6 days post-transduction, the percentage of GFP expressing cells was determined by flow cytometry analysis, as a readout of infectivity. To analyze the kinetics of integration, the condition that omitted DTG was set to 100% and this value was used to normalize earlier time points, for each virus independently. To compare the infectivity of the three viruses, the condition that omitted DTG in the PFV Gag WT transduction condition was arbitrarily set to 100%, and compared to the values of the same condition for the two other viruses.

For viral transduction of synchronized cells, 3.10^4^ HT1080 cells were platted in 24 well plates containing coverslips coated with 0.01% poly-L-Lysine (P4832, Sigma). The day after, HT1080 cells were treated with 20 µM of RO-3306 (SML0569, Sigma), in DMEM, for 17 h at 37°C. After 17 h of treatment, the medium was replaced by 500 µL of supernatant containing viral particles, supplemented with 20 µM of RO-3306, for 1 h at 37°C. After 1 h (after a total of 18 h of RO-3306 treatment), cells were carefully washed twice with warm fresh DMEM, and 1 mL of medium was added per well. This time point was considered as the starting point (T0) of the cell cycle analysis. Cells were fixed at different time points post RO-3306 release, 10 min at room temperature with 4% paraformaldehyde (PFA) (EM-15710, Euromedex) diluted in 1X PBS. Coverslips were kept in 1X PBS at 4°C, until further immunofluorescence analysis.

### Immunofluorescence

After fixation with 4% PFA, cells kept at 4°C were permeabilized for 5 min at room temperature (0.5% Triton X-100 diluted into 1X PBS) and washed twice with 1X PBS. Cells were incubated for 20 min at room temperature with IF Buffer (0.2% Triton X-100, 3% BSA diluted into 1X PBS). Primary antibodies were diluted in IF Buffer and incubated for 1 h at 37°C on cells. After two washes with 1X PBS, secondary antibodies diluted in IF Buffer were incubated on cells for 1 h at 37°C. Cells were then washed twice with 1X PBS and incubated with DAPI (D9542, Sigma) diluted 1:1000 in 1X PBS. Cells were then washed with successive baths of 1X PBS, H_2_O and absolute ethanol, coverslips were let air dried and a drop of mounting medium (Dako, Agilet) was added to seal coverslips on glass slides. Immunofluorescence analysis was performed by confocal microscopy, using a Leica TCS SP8 and the Leica LAS-X software, at the Bordeaux Imaging Center (BIC). The pinhole was set to 1, and z-stacks were collected at 0.5-µm intervals, with 7 planes collected per stack. Images were acquired at a 16-bit resolution with a pixel size of ∼90 nm. Images were analyzed using the Image J software.

### Image quantification

For image quantification, the Fiji software was used, and a semi-automated macro was designed to measure Gag-chromatin bound signal. Channels were split and analyses were performed on z-projections of 7 stacks. The cell outline was manually defined and a threshold was applied on the DAPI channel to define the chromatin region. Both the cell and chromatin areas were defined as region of interest (ROIs) for further analyses. A threshold on the Gag-channel was applied to select signals of interest. Finally, the area of Gag within the ROIs was measured, and a percentage of Gag-occupancy within these ROIs was automatically determined. Percentage were plotted on a graph, using Prism 9 software, representing the percentage of Gag signal measured within the chromatin area. A minimum of 5 cells per condition was analyzed. Means were represented with their standard deviation.

### Antibodies

For immunofluorescence analysis, the following antibodies were used: polyclonal rabbit antisera raised against PFV Gag recombinant^54^, dilution 1:1000; mouse anti-Lamin A/C (ab40567, Abcam), dilution 1:250; mouse anti-pericentrin (ab28144; abcam), dilution 1:500; mouse anti tubulin (T6199, Merck), dilution 1:500 and donkey cross-adsorbed secondary antibodies coupled with Alexa Fluor 488 or 594 (Life Technologies) were used at a dilution of 1:500. For Western blot analysis, the same polyclonal antisera raised against Gag was used at a dilution 1:1000 and HRP-conjugated anti-rabbit antibody (A8275, Sigma) was used at a dilution 1:10000.

### qPCR

1 h post-infection of HT1080 cells, DNA was extracted using the High Pure Viral Nucleic Acid Kit (Roche) according to the manufacturer’s instructions. A quantitative qPCR was performed to determine the relative quantity of viral DNA (vDNA), using the GoTaq® qPCR Master Mix (Promega). Primers used to detect vDNA were designed as follow^55^ : 203F – AGATTGTACGGGAGCTCTTCAC and 203R-CAGAAAGCATTGCAATCACC. The values obtained for DNA content of viral particle samples were expressed as percentage compared to the PFV Gag WT virus.

qPCR analysis was also performed to determine the relative quantification of vDNA integrated in HT1080 transduced cells. gDNA was extracted, using the Quick-DNA Miniprep Plus Kit (ZymoResearch). 120 ng of DNA was used per condition, and qPCR reactions were performed using the GoTaq® qPCR Master Mix. Primers 203F and 203F (described above) were used to detect vDNA, and GAPDH was used to normalized the gDNA quantity in each condition. The values obtained for vDNA detected per sample were expressed as percentage of the values obtained for a transduction with PFV Gag WT virus.

### Analyses of integration site distributions

PFV Integration sites for Y537Q mutant were generated for this study as previously described ^14^. WT PFV, R540Q variant and random integration control (RIC) sites were from accession numbers GSM1654885, <u>GSM2584470</u> and GSM2584468, respectively. We downloaded expression of mitotic genes from accession number GSE87476^35^, RepID ChIP-Seq data (GSM2123987) from the GSE80298 accession number^33^, and the hg38 coordinates of 10 SPIN states of the human genome from the published study^37^. Integration sites from the hg38 genome and SPIN coordinates were lifted to hg19 by the UCSC utilities LiftOver. Distributions of retroviral integration sites with respect to various genomic features were determined using BEDtools software suite^56^. Based on replication timing, human chromosomes were divided into three chromosomal groups: early (1, 15, 16, 17, 19, 20, 22), middle (3, 6, 7, 9, 10, 11, 12, 14,), and late (2, 4, 5, 8, 13, 18, 21) replicating chromosomes.

**Table 1.**
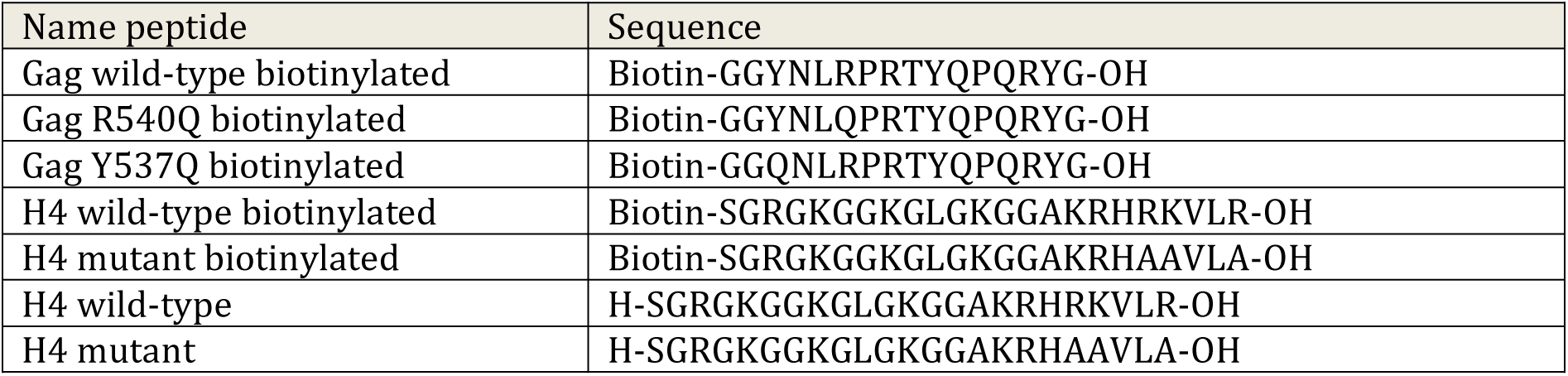
Sequences of peptides used for interaction experiments. Peptides were chemically produced.

---

**Figure S1.**
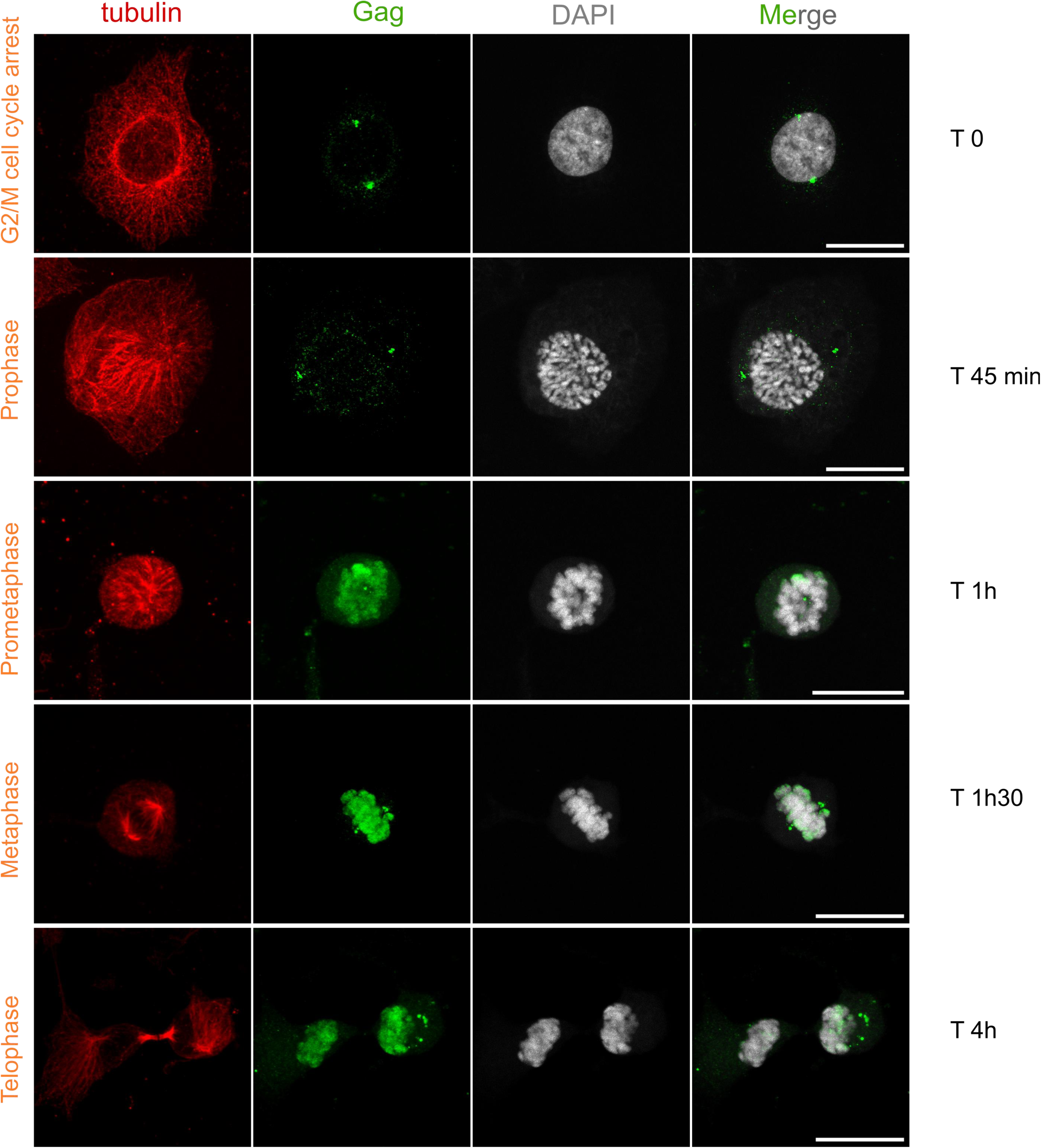
Optimization of cell cycle synchronization. HT1080 cells were treated with 20 µM RO-3306, transduced with PFV particles and fixed at different time points post RO-3306 release. Mitotic phases and their corresponding time points were identified via immunostaining of tubulin (red) and PFV Gag (green). Scale: 20 µm.

**Figure S2.**
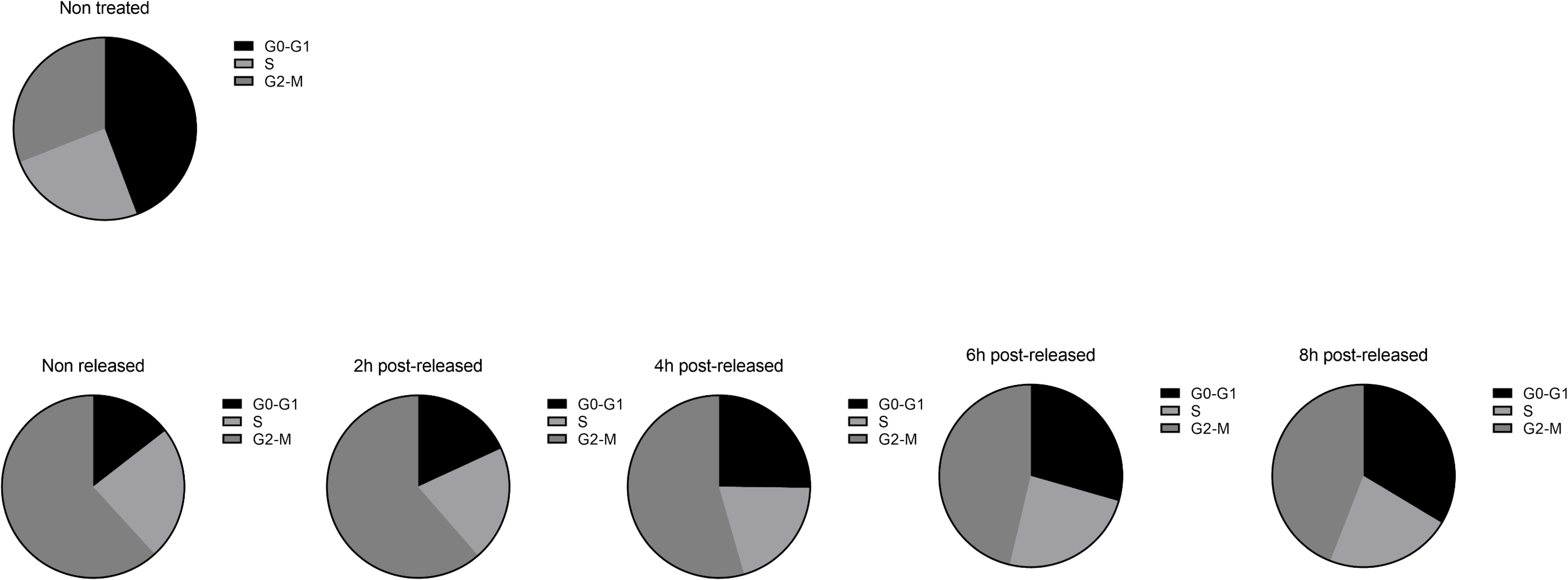
Cell cycle recovery of HT1080 cells after RO-3306 treatment. HT1080 cells were either treated with RO-3306 or not, and their DNA content was monitored at different time points using propidium iodide dye staining and FACS.

**Figure S3.**
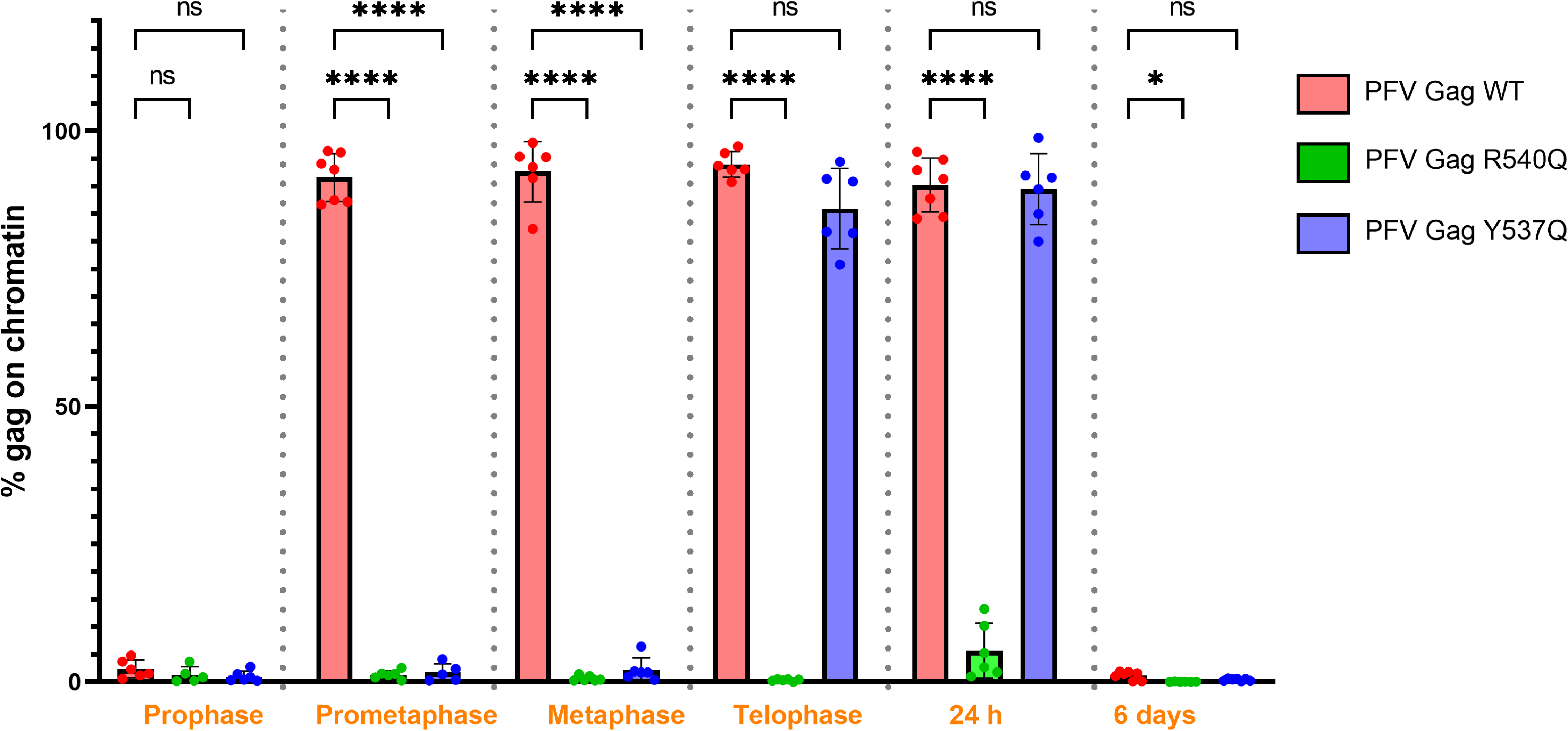
Quantification of Gag-chromatin binding as a function of mitotic phase. The percentage of chromatin surface recovered by Gag was quantified for each mitotic phase analyzed in Fig. 3. The mean of a minimum of 5 cells per condition was represented, with their corresponding standard deviation. Statistical analysis was performed using the two way ANOVA with the Tukey’s multiple comparisons tests (* p < 0.05; **** p < 0.0001).

**Figure S4.**
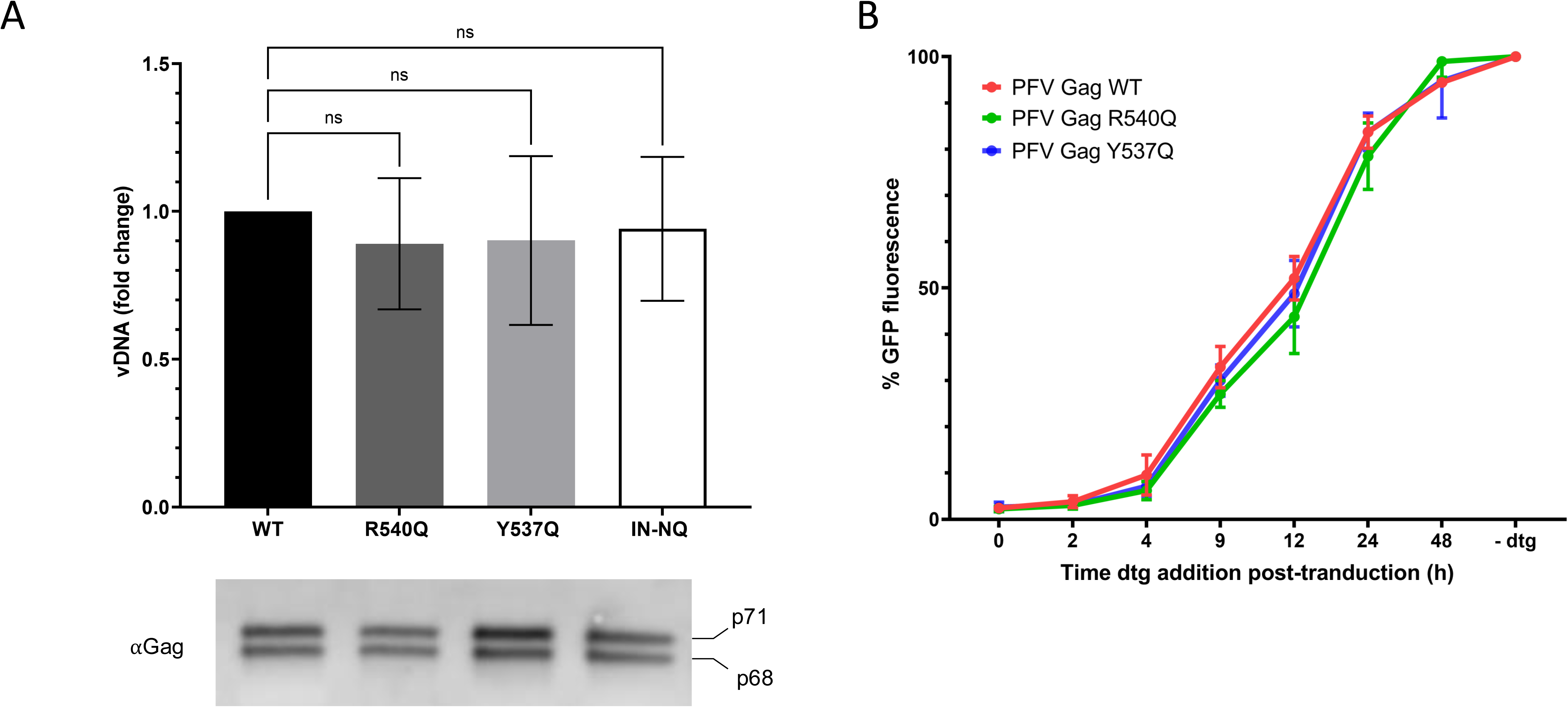
PFV particle loading controls. (A) Quantitative PCR of viral DNA used for transduction of HT1080 cells with PFV vector particles carrying WT, R540Q or Y537Q Gag with WT IN, or the WT Gag containing catalytically inert integrase (IN-NQ). Cells were harvested 1 h post-infection and the viral DNA was extracted and subjected to qPCR. The WT condition was set to 1 and results are expressed as comparative fold change. Immunoblots detecting Gag precursors (p71 and p68), which were used to control for viral input. The mean of three independent experiments is plotted. (B) Integration kinetics of WT and CBS mutants viruses monitored by inhibition of viral integration after dolutegravir (DTG) addition at different time points post-infection. Values without DTG for each condition were arbitrarily set to 100%. Relative means and standard deviations from three independent experiments are shown.

**Figure S5.**
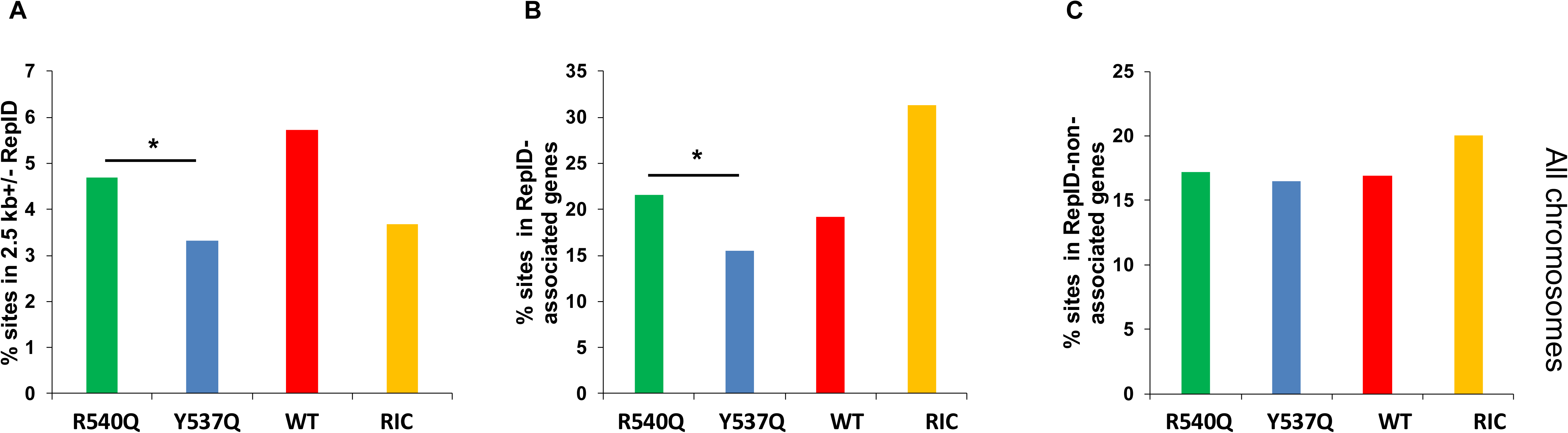
Correlation between replication-initiation determinant protein (RepID) binding sites and PFV integration. (A) PFV integration sites (%) within +/- 2.5 kb of RepID ChIP-Seq sites, (B) RepID-ChIP-Seq associated genes and (C) non-associated genes (p < 0.05; Fisher’s exact test).

**Figure S6.**
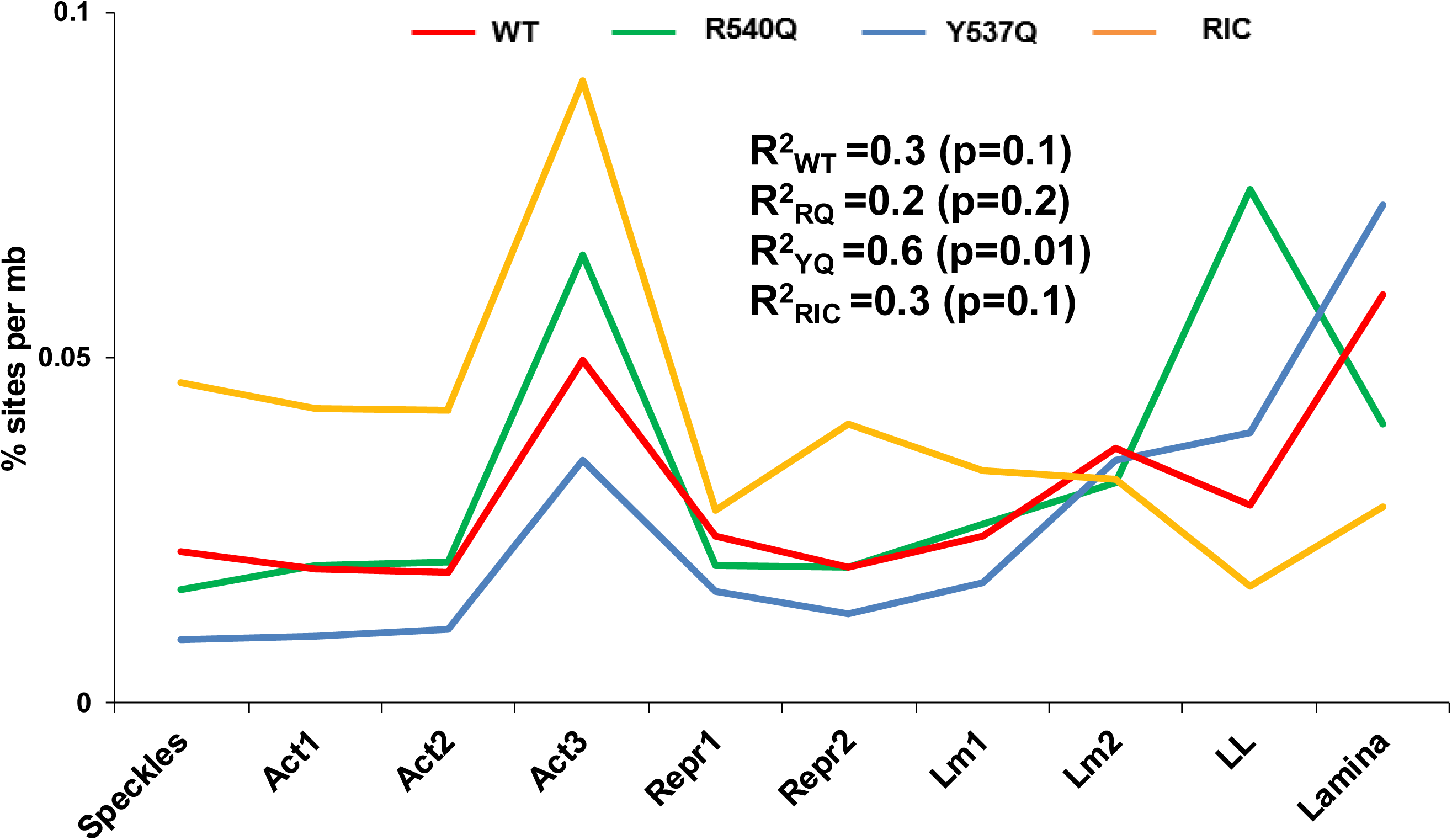
Distribution of PFV integration sites with respect to SPIN states. PFV integration sites (%) per Mb across 10 Spatial Position Inference of the Nuclear genome SPIN states (X-axis) were calculated for WT, R540Q (RQ) and Y537Q (YQ) PFVs along with a random integration control (RIC). Coefficients of determination (R^2^) between integration % (Y-axis) and SPIN states are shown (calculated in Excel and square root of R^2^ was used to calculate p values at https://www.socscistatistics.com/pvalues/pearsondistribution.aspx). Act1, Interior Active 1; Act2, Interior Active 2; Act3, Interior Active 3; Repr1, Interior Repressive 1; Repr2, Interior Repressive 2; Lm1, Near Lamina 1; Lm2, Near Lamina 2; LL, Lamina-Like.

**Figure S7.**
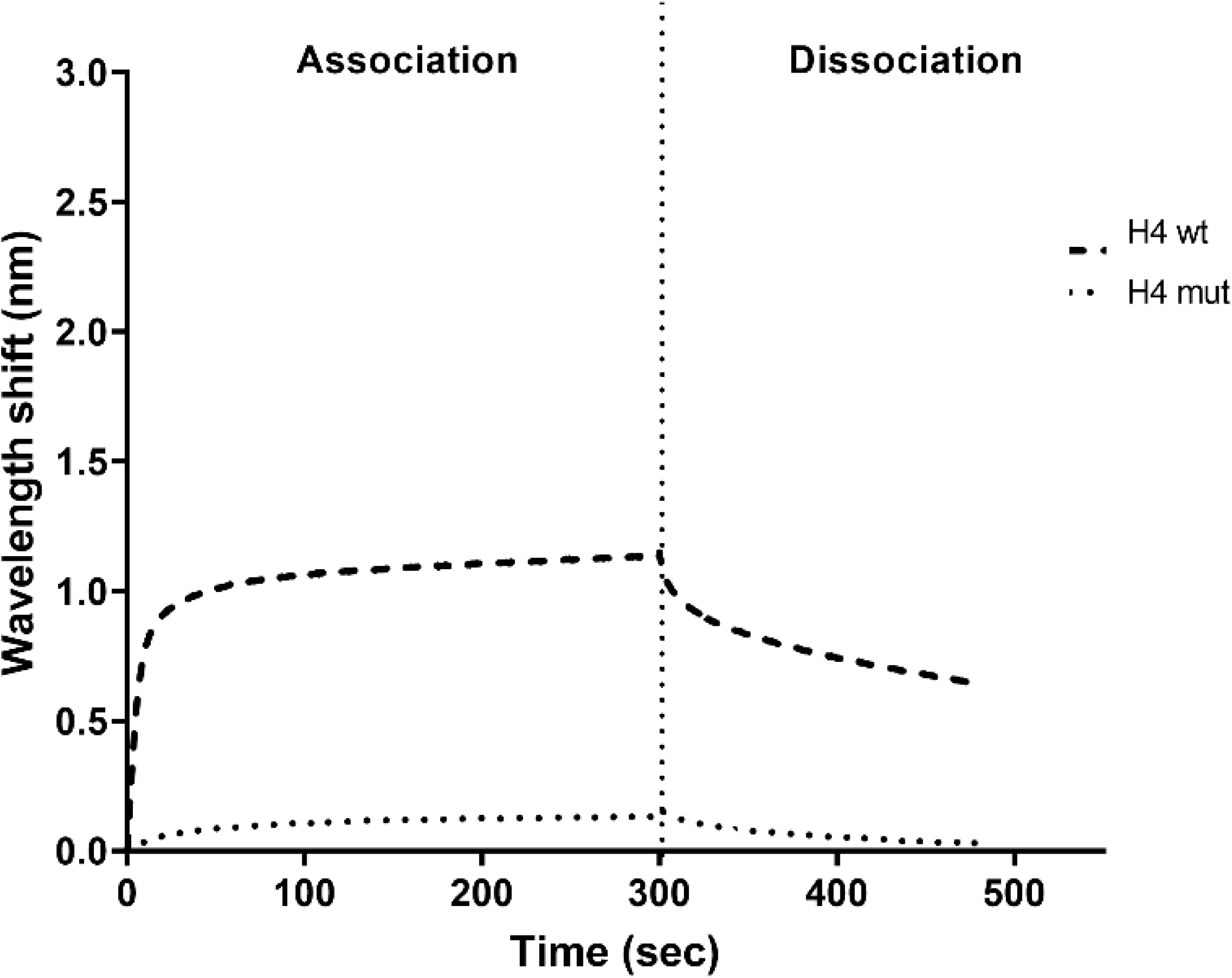
H4 tail-NCP interaction controls. Bio-Layer interferometry (BLI) sensorgram of free nucleosomes binding to immobilized WT H4 tail (residues 2-24) (dashed line) or H4 mut (triple alanine substitution) (dotted line). The binding intensity (nm) is normalized with a condition without NCP. The mean of 2 independent experiments is plotted.

